# Polygenic Risk for Alcohol Misuse is Moderated by Romantic Partnerships: Primarily in Men

**DOI:** 10.1101/528844

**Authors:** Peter B. Bar, Sally I-Chun Kuo, Fazil Aliev, Antti Latvala, Richard Viken, Richard J. Rose, Jaakko Kaprio, Jessica E. Salvatore, Danielle M. Dick

**Author notes:** joint last authors (Jessica E. Salvatore, Danielle M. Dick). Corresponding Authors: Peter Barr, PhD, Department of Psychology, Virginia Commonwealth University, 8 North Harrison St., Richmond, VA. 23284, Danielle M. Dick, PhD, Department of Psychology, Virginia Commonwealth University, 800 W. Franklin Street, Box 842018, Richmond, VA 23284-2018, (804)828-8756.

## Abstract

**Importance:** Problematic alcohol use remains a leading influence on preventable mortality and morbidity across the globe. Those in committed relationships consistently report lower levels of alcohol misuse and problems.

**Objective:** To determine 1) whether genetic risk for alcohol misuse is moderated by romantic relationships (gene-environment interaction; GxE), and 2) whether GxE results are consistent across sex.

**Design:** Data came from the young adult wave of the Finnish Twin Study (FinnTwin12), a nationally representative sample of twins. Predictors included genome-wide polygenic scores (GPS), derived from a recent genome-wide association study (GWAS) of alcohol consumption in ~1 million participants; and participant reports of relationship status.

**Setting:** Finland

**Participants:** An intensively studied subset of FinnTwin12 received a diagnostic interview during the young adult phase (1,312 of 1,347 individuals provided genotypic data). The analytic sample includes those with complete interview and genetic data (N=1,201, 54% female).

**Exposure:** Self-reported involvement in a romantic partnership.

**Main Outcomes and Measures:** Drinking frequency, intoxication frequency, and DSM-IV alcohol dependence (AD) symptoms from a diagnostic interview.

**Results:** GPS predicted drinking frequency (b = 0.109; 95% CI = 0.051, 0.167), intoxication frequency (b = 0.111; 95% CI = 0.054, 0.168), and AD symptoms (b = 0.123; 95% CI = 0.064, 0.182). Relationship moderated the association between GPS and drinking frequency (b = −0.105; 95% CI = −0.211, −0.001), intoxication frequency (b = −0.118; 95% CI = −0.220, −0.015), and AD symptoms (b = −0.119; 95% CI = −0.229, −0.010). The interaction for drinking frequency was not significant after correcting for covariates. There was a 3-way interaction between sex, relationship status, and GPS for intoxication frequency (b = 0.223; 95% CI = 0.014, 0.432), with the two-way interaction of relationship status and PRS on intoxication frequency being significant only in men.

**Conclusions and Relevance:** Being in a relationship reduced the association between genetic predisposition and high risk drinking. Part of the protective effect of committed partnerships on alcohol misuse observed in epidemiological research may be in limiting genetic liability. However, this protective effect was largely limited to males, mapping onto earlier findings suggesting that males benefit more from romantic partnerships.

**Key Points:** *Question:* Do romantic relationships moderate polygenic risk on alcohol misuse in young adulthood?

*Findings:* Involvement in romantic relationships moderated the polygenic risk on frequency of intoxication and DSM-IV alcohol dependence symptoms, such that polygenic associations with alcohol misuse were stronger among those not in a romantic relationship. Males experienced a stronger protective effect of romantic relationship in limiting the manifestation of genetic predispositions toward intoxication frequency.

*Meaning:* The interplay between genes and environment is important in understanding etiology of problematic alcohol use, and romantic relationships appear to buffer genetic risk for alcohol misuse in young adulthood. Findings underscore how social relationships may alter the risk posed by genetic predispositions.

---

Alcohol use is one of the leading contributors to preventable mortality and morbidity, worldwide.^1–3^ Twin and family studies indicate that genetic influences account for approximately 50% of the variation in the population^4^; however, there is strong evidence the importance of genetic influences changes across environmental contexts (gene-environment interaction, or G×E).^5,6^ Environments that allow greater access to alcohol, or acceptance of alcohol use, may create opportunity for increased manifestation of individual predispositions toward alcohol misuse and consequently the development of problems.^7–11^ Conversely, environments that exert more social control, such as greater parental monitoring in adolescence, appear to reduce the importance of genetic predispositions.^7,12^ Mapping which environments reduce alcohol misuse among those at greater genetic risk will be critical for developing tailored prevention intervention strategies as we move into an era of precision medicine.

Much of the foundational work on G×E in alcohol outcomes has been conducted in twin studies.^6–9,12^ Most G×E studies to date using measured genotypes on alcohol use outcomes have focused on candidate genes or single nucleotide polymorphisms (SNP), where the effect of a specific candidate gene or single SNP varies as a function of the environment.^6^ However, candidate gene research has generated inconsistent results, likely a reflection of being underpowered to robustly detect moderations, false positives, and publication bias.^13,14^ Furthermore, the use of single genes in G×E studies does not align with our current molecular genetic understanding that complex behaviors, including alcohol use,^15^ problems,^16^ and dependence,^17^ have a polygenic architecture, driven by many genetic variants of very small effect.^18,19^ Large sample sizes are needed to detect robust genetic associations for complex behavioral outcomes in genome-wide association studies (GWAS), which use data from the entire genome rather than relying on predefined SNPs.^20,21^

To characterize individual risk across hundreds or thousands of alleles associated with an outcome in a GWAS, genome-wide polygenic risk scores (GPS) have emerged as a way to aggregate this information into a single score. As we begin to identify GPS robustly associated with substance use and dependence, one of the critical next steps toward precision medicine will be to characterize the pathways by which risk unfolds.^22^ For alcohol-related outcomes, this will necessitate characterizing how specific environments moderate the likelihood that individuals carrying risky genetic predispositions will develop excessive use, problems, and dependence, providing important information about targeted areas for intervention.

In this study, we focused on romantic relationships, as epidemiological research has consistently shown that those in committed relationships (especially marriage) engage in fewer risky or health-deteriorating behaviors, such as alcohol misuse.^23,24^ This reduction in risky behaviors is due in part to increased social control and monitoring associated with being in a relationship,^23^ as well as individuals’ motivation to align their behavior with the social expectations typically associated with the spousal role.^25,26^ However, marriage-like relationships are generally beneficial for men but more or less indifferent for women,^27^ suggesting important sex-differences in any protective effect. Finally, twin studies have found that the heritability of alcohol consumption is decreased among individuals in committed relationships,^28,29^ suggesting that being with a partner may act as a “social control” that limits expression of genetic predispositions toward alcohol problems.

Here, we test this hypothesis using molecular genetic data in a population-based sample of young adults.^30^ We focused on young adulthood because it is a critical period for the development of alcohol use patterns and problems,^28^ with heavy alcohol use at its highest point^31^ and the peak age of onset for alcohol related disorders falling during this period.^32^ We used results from the largest mega-analysis to date on alcohol consumption (drinks per week in ~1 million individuals),^15^ to calculate genome-wide polygenic scores in our independent, population-based sample. We tested the association of these polygenic risk scores with alcohol use, heavy consumption, and alcohol problems, and importantly, whether being in a romantic relationship changed the association between genetic risk and alcohol outcomes. Finally, because there are sex differences in patterns of alcohol use and in the prevalence of alcohol use disorders^32^ and heavy consumption,^31^ we examined whether there were sex differences in G×E.^33^

## Methods

### Sample

Data come from the youngest cohort of the Finnish Twin Cohort Study (FinnTwin12). Families were identified from Finland’s Population Registry, permitting comprehensive nationwide ascertainment for twins born from 1983 to 1987. Baseline collection occurred when twins were approximately 12 years old, with a sample of approximately 5600 twins (87% participation) and their families.^30^ Follow-up surveys occurred at ages 14, 17.5, and during young adulthood (age range 20-26). Twin zygosity was determined using items developed for twin children.^34^ Confirmation by multiple genetic markers revealed that 97% of same-sex pairs retained the original questionnaire-based zygosity classification^35^. The Helsinki University Central Hospital District’s Ethical Committee and Indiana University’s Institutional Review Board approved the FinnTwin12 study. Of those in the larger sample, a subset of intensively studied individuals also received in-depth clinical interviews (N = 1,347) and participated in DNA collection as young adults. In the present study, we limited our analyses to those who had complete information on all relevant study variables and who had initiated alcohol use (n = 1,201). The analytic subset did not differ significantly from the full sample in terms of demographic characteristics or alcohol misuse (see supplemental information for more detail).

### Genotyping and Quality Control

Genotyping was conducted using the Human670-QuadCustom Illumina BeadChip at the Wellcome Trust Sanger Institute.^36^ Quality control steps included removing SNPs with minor allele frequency (MAF) <1%, genotyping success rate <95%, or Hardy–Weinberg equilibrium *p* < 1 × 10^−6^, and removing individuals with genotyping success rate <95%, a mismatch between phenotypic and genotypic gender, excess relatedness (outside of known families), and heterozygosity outliers. Genotypes were imputed to the 1,000 Genomes Phase reference panel^37^ reference panel using ShapeIT^38^ for phasing and IMPUTE2^39^ for imputation, resulting in 13,688,418 autosomal SNPs for analyses. Prior analyses indicated a single dimension of ancestry in the sample.^40^ Although a single dimension of ancestry does not preclude variation along this dimension, we note that fine-scale population substructure is less of an issue for common variants (vs. rare variants), especially in the present sample given the relatively longer LD blocks that make the Finnish population more homogenous than other populations of mixed European ancestry.

### Measures

*Alcohol-Related Behaviors* were assessed across increasing levels of severity. Drinking frequency was measured by asking “How often do you use alcohol?” Responses included “never” (0), “once a year” (1), 2-4 times a year (2), “every other month” (3), “once a month” (4), “more than once a month” (5), “once a week” (6), “more than once a week” (7), and “daily” (8). Intoxication frequency was assessed by asking “How often do you use alcohol in such a way that you get really drunk?” Responses were the same for drinking frequency. We transformed these ordinal measures into pseudo-continuous measures of the frequency of these behaviors in the past 30 days.^41,42^ Finally, we included a count of lifetime DSM-IV Alcohol Dependence (AD) symptoms, assessed using the Semi-Structured Assessment for the Genetics of Alcoholism (SSAGA), a reliable and valid clinical instrument.^43^ Each alcohol measure was log transformed (left anchored at 1) to adjust for positive skew. *Relationship Status* was measured by asking, “How long (in years) have you been together with your present partner?” Respondents that indicated they were not in a relationship were coded as 0. Those who indicated they were in a romantic relationship for any length were coded as 1. We ran sensitivity analyses with a stricter definition of relationship status (those in a relationship >= 2 years). Our results did not fundamentally differ from the more inclusive definition and we retained the original measurement of relationship status. Finally, we included age, sex, educational attainment (based on the Finnish education system: basic education; vocational training; secondary education; tertiary education), and whether or not respondents were still in school^44^ (dichotomous yes/no) as covariates.

### Genome-wide Polygenic Scores

We created polygenic scores derived from a large-scale GWAS of number of alcoholic drinks per week in approximately one million individuals^15^ provided by the GWAS & Sequencing Consortium of Alcohol and Nicotine Use (GSCAN). As FinnTwin12 was included in the original discovery GWAS, we obtained summary statistics with all Finnish participants, including FinnTwin12 participants removed. There were 3,707,235 autosomal SNPs in common after QC. We used the well-established process of clumping and thresholding.^45^ SNPs from the discovery GWAS were clumped based on linkage disequilibrium (LD) using the *clump* procedure in PLINK,^46^ based on an R^2^ = .25, with a 500kb window, resulting in 407,604 independent SNPs for creating scores. We then created scores based on differing thresholds of GWAS p-values (p<.0001, p<.001, p<.01, p<.05, p<.10, p<.20, p<.30, p<.40, p<.50). We converted GPS to Z-scores for interpretation.

We note that alcohol consumption and problematic use, though highly correlated, have distinct genetic influences.^47^ We ran a series of sensitivity analyses to determine if recent GWAS focused on alcohol problems or dependence^16,17^ provided better assessments of genetic liability for alcohol misuse (see supplemental information). However, in each case, the scores derived from GSCAN were the most predictive.

### Analytic Strategy

First, we estimated the effect of GPS across each p-value threshold to determine the most predictive score (based on model R^2^) for each alcohol phenotype. We then tested whether relationship status moderated the association of the genome-wide polygenic scores. In the instances where we found evidence for a significant interaction, we fit a more robust model for evaluating G×E,^48^ which includes all G × covariate and E × covariate interaction terms. Finally, we tested for sex-specific G×E by including a three-way interaction term. We determined whether estimates were significant using an *α* of p < .05 (two-sided test). Because the FinnTwin12 data is a family-based data set, we evaluated all hypotheses using a linear mixed model with random intercepts for each family in the *lme4*^49^ package in in *R* 3.5.1.^50^ We estimate effect size (R^2^) using a method designed for mixed effects models^51^ with the *MuMIn* package.^52^

## Results

Males exhibited higher mean levels of each alcohol measure (Table 1). The alcohol phenotypes were also modestly correlated (*r*_drinking*intox_ = .64, *r*_drinking*ADsx_ = .37, *r*_intox*ADsx_= 43), with stronger correlations between the consumption items than with the measure of AD symptoms.^47^

**Table 1:** Descriptive Statistics for Finnish Twin Study (FinnTwin12)

| | Males (N = 551) | | Females (N = 650) | | $\chi^2$ / t-test |
| --- | --- | --- | --- | --- | --- |
|  | Mean/N | SD/% | Mean/N | SD/% |  |
| Drinking Frequency | 5.10 | 4.54 | 3.43 | 3.24 | * |
| Intoxication Frequency | 2.02 | 1.97 | 1.12 | 1.44 | * |
| DSM-IV Alcohol Dependence Symptoms | 1.54 | 1.35 | 1.29 | 1.37 | * |
| GPS<sup>†</sup> | -0.03 | 1.02 | 0.03 | 0.98 |  |
| Age | 21.94 | 0.77 | 21.95 | 0.76 |  |
| Educational Attainment |  |  |  |  | * |
| Basic Education | 38 | 6.9% | 30 | 4.6% |  |
| Vocational Training | 207 | 37.6% | 157 | 24.2% |  |
| Secondary Education | 299 | 54.3% | 424 | 65.2% |  |
| Tertiary Education | 7 | 1.3% | 39 | 6.0% |  |
| Enrolled in school | 285 | 51.7% | 401 | 61.7% | * |
| In relationship | 269 | 48.8% | 416 | 64.0% | * |

\*p < .05 for Chi-square/T-test difference between males and females
<sup>†</sup> Standardized (Z-scores) GPS including SNPs with p < .50

### Polygenic Score Performance

Figure 1 provides the incremental R-squared for polygenic scores at different p-value inclusion thresholds. The variance explained at each p-value threshold in GPS represents the change in R-squared from the baseline model (age and sex as covariates) after including the GPS at that p-value threshold. GPS were significantly associated with each alcohol related behavior across almost all of the p-value thresholds, with the exception of the most restrictive scores in relation to drinking frequency. GPS explained more variance as p-value thresholds became more inclusive, peaking and leveling off at thresholds between p < .20 and p < 0.50. We decided to use the most liberal threshold (p < .50) for all models going forward.

**Figure 1:**
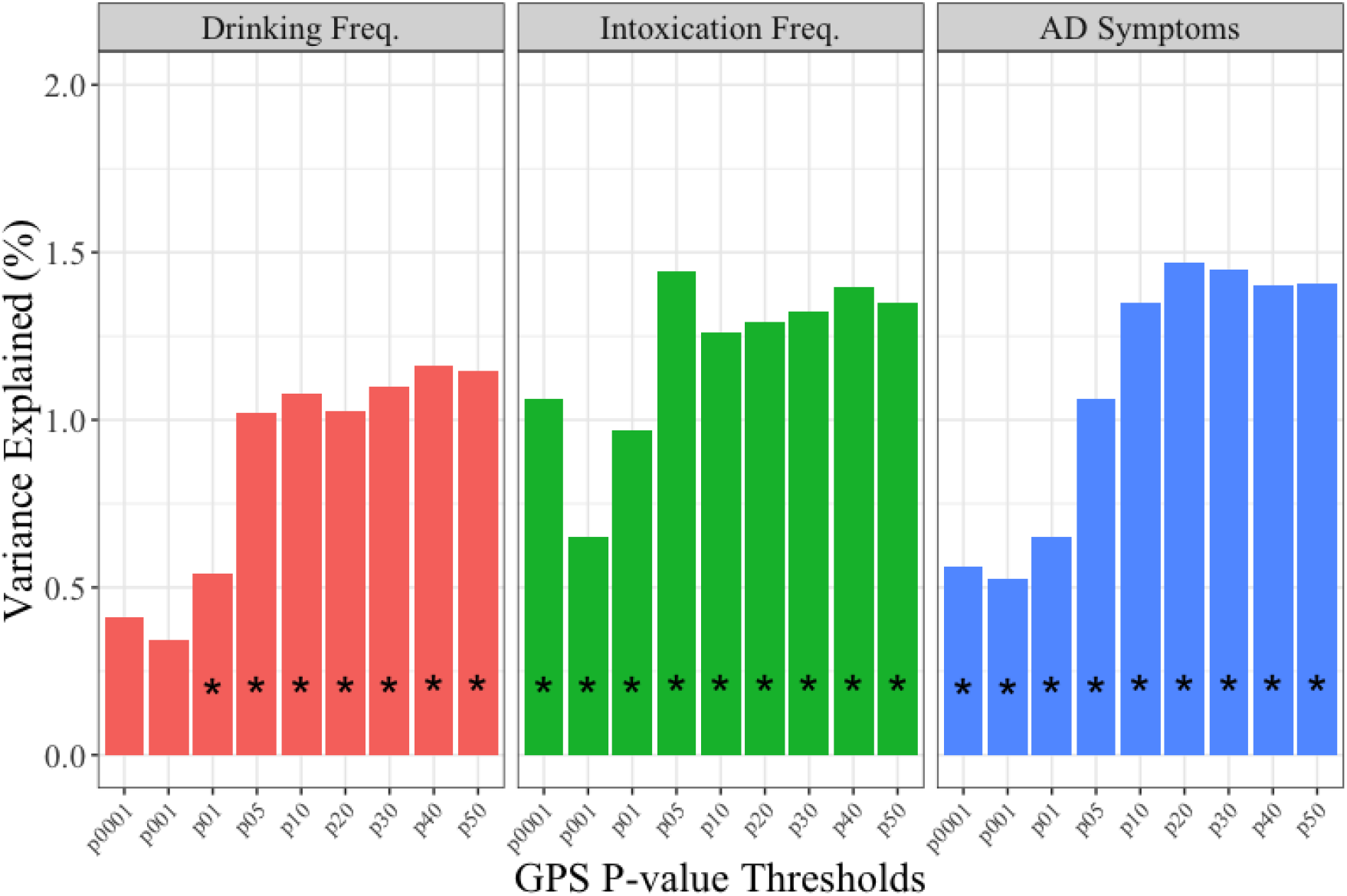
Predictive Power of GSCAN Polygenic Scores. Change in model R2 from base model (age and sex as covariates) to model including polygenic scores at various p-value inclusion thresholds (determined by p-value from discovery GWAS). * association p < .05.

In order to ensure the GPS were predictive of alcohol problems above and beyond levels of consumption (which are genetically correlated but distinct phenotypes),^47^ we estimated the effect of GPS while accounting for either drinking or intoxication frequency. GPS were significantly related to AD symptoms after statistically controlling for drinking frequency (b = 0.085, p < .01) or intoxication frequency (b = 0.075, p < .01; see supplemental information for full results). Finally, we estimated the polyserial correlation between GPS and relationship status (ρ = 0.005, p > .05) to assess the possibility of gene-environment correlation.

### Main Effects of Polygenic Score and Relationship Status

Table 2 provides the estimates for the linear mixed models evaluating the joint effect of GPS and relationship status. In the model for main effects (Model 1), those currently in a relationship had lower levels of intoxication frequency, but not drinking frequency or AD symptoms. GPS remained significantly associated with each of these alcohol related behaviors.

**Table 2:** Linear Mixed Models for Alcohol Related Behaviors (N = 1,201)

|  | <u>Model 1</u> |  |  | <u>Model 2</u> |  |  | <u>Model 3</u> |  |  |
| --- | --- | --- | --- | --- | --- | --- | --- | --- | --- |
|  | B | SE |  | B | SE |  | B | SE |  |
| <u>Drinking Frequency</u> |  |  |  |  |  |  |  |  |  |
| Female | -0.456 | 0.059 | *** | -0.458 | 0.059 | *** | -0.437 | 0.086 | *** |
| In relationship | -0.090 | 0.055 |  | -0.087 | 0.055 |  | -0.069 | 0.079 |  |
| GPS | 0.109 | 0.030 | *** | 0.169 | 0.043 | *** | 0.193 | 0.055 | ** |
| In relationship*GPS | - | - | - | -0.105 | 0.054 | * | -0.158 | 0.076 | * |
| Female*GPS | - | - | - | - | - | - | -0.061 | 0.086 |  |
| Female*In relationship | - | - | - | - | - | - | -0.038 | 0.110 |  |
| Female*In Relationship*GPS | - | - | - | - | - | - | 0.109 | 0.110 |  |
| Pseudo-R<sup>2</sup> | 0.073 |  |  | 0.076 |  |  | 0.077 |  |  |
| <u>Intoxication Frequency</u> |  |  |  |  |  |  |  |  |  |
| Female | -0.543 | 0.058 | *** | -0.544 | 0.058 | *** | -0.535 | 0.084 | *** |
| In relationship | -0.178 | 0.054 | ** | -0.176 | 0.054 | ** | -0.171 | 0.077 | * |
| GPS | 0.111 | 0.029 | *** | 0.179 | 0.042 | *** | 0.239 | 0.054 | *** |
| In relationship*GPS | - | - | - | -0.118 | 0.052 | * | -0.222 | 0.073 | ** |
| Female*GPS | - | - | - | - | - | - | -0.149 | 0.084 |  |
| Female*In relationship | - | - | - | - | - | - | -0.016 | 0.107 |  |
| Female*In Relationship*GPS | - | - | - | - | - | - | 0.223 | 0.107 | * |
| Pseudo-R<sup>2</sup> | 0.110 |  |  | 0.114 |  |  | 0.117 |  |  |
| <u>AD Symptoms</u> |  |  |  |  |  |  |  |  |  |
| Female | -0.197 | 0.061 | ** | -0.199 | 0.061 | ** | -0.123 | 0.089 |  |
| In relationship | -0.097 | 0.057 |  | -0.095 | 0.057 |  | -0.028 | 0.082 |  |
| GPS | 0.123 | 0.030 | *** | 0.191 | 0.044 | *** | 0.196 | 0.057 | ** |
| In relationship*GPS | - | - | - | -0.119 | 0.056 | * | -0.154 | 0.079 |  |
| Female*GPS | - | - | - | - | - | - | -0.018 | 0.089 |  |
| Female*In relationship | - | - | - | - | - | - | -0.134 | 0.115 |  |
| Female*In Relationship*GPS | - | - | - | - | - | - | 0.069 | 0.114 |  |
| Pseudo-R<sup>2</sup> | 0.029 |  |  | 0.032 |  |  | 0.034 |  |  |

All models include age, educational attainment, and student status as covariates. Family clustering adjusted for by including random intercepts for the family level. GPS and alcohol phenotypes were standardized.
\* p < .05; \*\* p < .01; \*\*\* p < .001

### Gene-Environment Interaction Models

Model 2 (Table 2) presents the estimates for G×E. There was a significant interaction between relationship status and polygenic scores for each alcohol behavior. We refit each of these models with interactions between relationship status and each covariate and interactions between GPS and each covariate (plotted in Figure 2, see supplemental information for full results) to account for possible confounding.^48^ P-values were attenuated, especially in the models for drinking frequency and AD symptoms, but the nature of the interactions remained unchanged for the other phenotypes. The shape of the interaction was similar across all phenotypes, but most pronounced for intoxication. In the case of intoxication frequency, there was a stronger association between genetic risk score and intoxication frequency among individuals who are not in romantic relationships, and a relatively weaker association between genetic risk score and intoxication frequency among those who were in romantic relationships.

**Figure 2:**
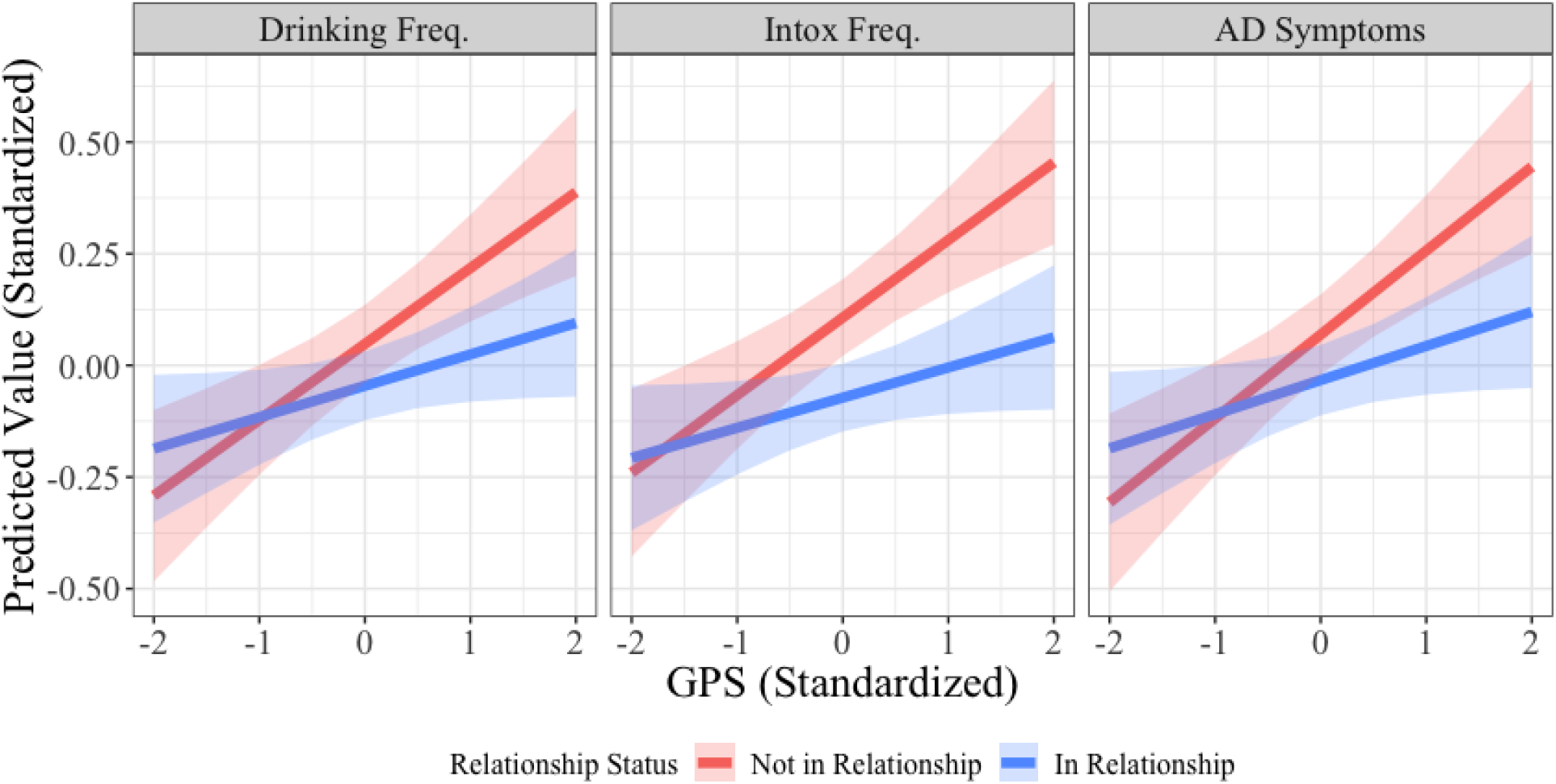
Gene-Environment Interaction across Relationship Status and Polygenic Risk. Predicted values of each alcohol phenotype (standardized) across the range of polygenic scores for those in a relationship (blue) and those not in a relationship (red). Shaded areas represent 95% pointwise confidence intervals of estimates.

### Sex Differences in G×E

Finally, we tested for sex differences in the interaction between relationship status and GPS. We found no evidence of a significant three-way interaction between sex, relationship status, and GPS for either drinking frequency or AD symptoms. However, we did find a significant three-way interaction in the models for intoxication frequency. This interaction remained significant even after adjusting for possible confounding in the G×E interactions. Figure 3 displays the predicted values from this model. For intoxication frequency, the G×E effect appears to be driven by the effect in males.

**Figure 3:**
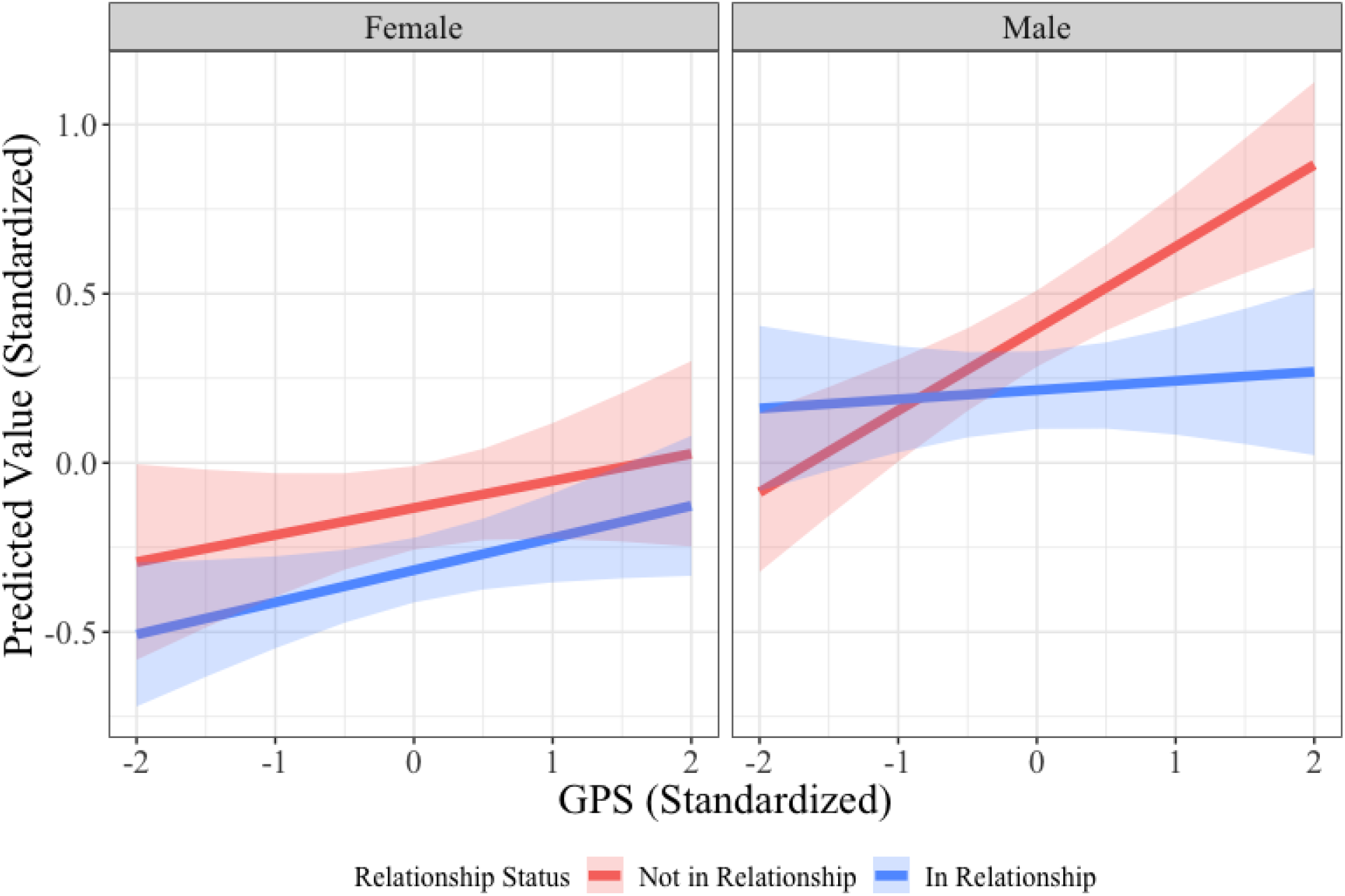
Sex Differences in G×E for Intoxication Frequency. Predicted values of intoxication frequency (standardized) across the range of polygenic scores and sex for those in a relationship (blue) and those not in a relationship (red). Shaded areas represent 95% pointwise confidence intervals of estimates.

## Discussion

We tested whether polygenic risk scores derived from a meta-analysis of alcohol consumption were associated with alcohol outcomes in an independent, population-based young adult sample, whether romantic relationship status moderated the association of genetic predispositions with alcohol outcomes, and whether observed effects varied between females and males.

Polygenic scores derived from variants associated with consumption are predictive of use, misuse, and problems among young adults. As hypothesized, being in a romantic relationship moderated the association between GPS and each alcohol phenotype (drinking frequency, intoxication frequency and AD symptoms). Similar to previous twin research,^53,54^ among individuals with elevated genetic predisposition, levels of misuse were lower in those in a romantic partnership. We posit that the constraints and responsibilities placed on individuals within romantic partnerships limits their ability to express underlying predispositions towards alcohol misuse, fitting with the social control model of gene-environment interaction.^23,55^ Additional inspection (available in supplemental information) revealed these interactions did not appear to be driven by outliers at either end of the distribution.

Finally, we examined whether there were sex differences in these G×E effects. We found no evidence of sex differences in the G×E effect for drinking frequency or AD symptoms. However, the G×E effect for intoxication frequency was driven primarily by the effect in males. Previous work in social epidemiology has documented how males tend to “over-benefit” from relationships in terms of health.^27^ This may reflect the tendency for women in relationships to be the emotional and social support providers, of which men are the receivers.^56^ In the current study, we see that this effect may be due in part to limiting genetic liability among a riskier drinking group (see supplemental information for additional analyses). This difference does not appear in AD symptoms, which is likely the result of these symptoms measuring aspects of both consumption and problems. Any role relationship status has in limiting genetic liability seems limited to levels of heavy consumption.

Our findings have important practical implications for researchers and clinicians interested in those at greater risk for alcohol misuse. First, the signal for genetic associations may be drastically reduced in young adults in a committed relationship. Future research on gene identification efforts may benefit from the inclusion of important environmental information in order to increase power to detect genetic variants associated with various forms of alcohol misuse. Considering G×E in the discovery GWAS may be of even more importance in regards to alcohol use phenotypes, as there is consistent evidence of G×E from twin studies.^41,53,57,58^ For clinicians, these analyses point to committed relationships as a malleable environmental condition that may help reduce individuals’ level of misuse, in part, by limiting realization of genetic predisposition. Although gene-environment correlation is always an important consideration, we note that our GPS was uncorrelated with relationship status, consistent with previous research using more causally-identified designs.^59^

This research has several limitations. First, although the polygenic scores explained more variance in these outcomes than previous iterations using smaller discovery GWAS, the variance explained by the largest meta-analysis of alcohol consumption to date, compiling data from ~1 million individuals, continued to be small (R^2^ ~ .015), especially compared to other complex phenotypes with similar sample sizes, like education attainment (R^2^ ~ .12)^60^. Discovery samples with better phenotyping will be necessary to create scores that explain the total SNP-based heritability. Second, though we found evidence of G×E, it does not rule out other confounding factors. Larger twin samples with genotypic data that allow for within-family designs will help to further account for possible environmental confounders shared across families (e.g. neighborhood factors, religiosity, socioeconomic status; see supplemental for sensitivity analyses). Third, our measure of romantic partnerships did not examine which relationship characteristics moderate polygenic scores (e.g. relationship quality, partner’s drinking, emotional support). Finally, our measure of AD symptoms was a lifetime measure. Supplemental analyses revealed similar patterns between lifetime and past 12-month symptoms.

In conclusion, polygenic scores from a large-scale GWAS of drinks per week predicted levels of alcohol use and misuse among a sample of young adults. However, the likelihood an individual carrying riskier genetic predispositions would display problematic patterns of use changed as a function of the environment. Individuals at greater genetic risk who were in romantic relationships were less likely to misuse alcohol. For drinking to intoxication, this interaction appears to occur primarily among males. This finding is consistent with previous research on social determinants of health that men tend to over-benefit from romantic partnerships.^27^ This research underscores the importance of considering the interplay between genes and environment when considering etiology and intervention for problematic alcohol use. In order for genetic risk scores to be useful in clinical settings, we must understand how genetic risk interacts with the environment.

## Acknowledgments

Research reported in this publication was supported by the National Institute on Alcohol Abuse and Alcoholism of the National Institutes of Health under award numbers R01AA015416, K02AA018755, K01AA024152, and F32AA022269; the Academy of Finland (grants 100499, 205585, 118555, 141054, 265240, 263278, and 264146); and the Scientific and Technological Research Council of Turkey (TÜBİTAK) under award number 114C117. The content is solely the responsibility of the authors and does not necessarily represent the official views of the National Institutes of Health, the Academy of Finland, or the Scientific and Technological Research Council of Turkey. The authors have no conflict of interests to report.

