## Supplemental info for "Polygenic Risk for Alcohol Misuse is Moderated by Romantic Partnerships: Primarily in Men"

**Comparison of Full and Analytic Sample**

Of the original 1,347 individuals who participated in the young adult diagnostic interview, 1,286 also completed a survey that included questions covering a broad array of areas (including romantic partnership and alcohol consumption). Of this group, 1,247 also provided genotypic data (including those inferred from MZ co-twins). Amongst this final subset, 32 individuals had not initiated alcohol use and another 14 had missing data on one of the items resulting in a final sample size of 1,201. The table below provides the descriptive statistics from all who answered a question compared to those included in the analyses. Overall we see no observed differences among the full and reduced samples.

|  | **Full Sample** | | | **Analytic Sample** | | |
| --- | --- | --- | --- | --- | --- | --- |
|  | Valid N | Mean/N | SD/% | Valid N | Mean/N | SD/% |
| Female | 1321 | 694 | 52.5% | 1201 | 650 | 54.1% |
| Age | 1347 | 21.95 | 0.77 | 1201 | 21.94 | 0.77 |
| Drinking Frequency | 1252 | 4.22 | 4.07 | 1201 | 4.20 | 3.98` |
| Intoxication Frequency | 1252 | 1.53 | 1.76 | 1201 | 1.54 | 1.76 |
| DSM-IV Alcohol Dependence Symptoms | 1253 | 1.40 | 1.37 | 1201 | 1.40 | 1.37 |
| Educational Attainment: |  |  |  |  |  |  |
| Basic Education | 1282 | 73 | 5.7% | 1201 | 68 | 5.7% |
| Vocational Training | 1282 | 395 | 30.8% | 1201 | 364 | 30.3% |
| Secondary Education | 1282 | 763 | 59.5% | 1201 | 723 | 60.2% |
| Tertiary Education | 1282 | 51 | 4.0% | 1201 | 46 | 3.8% |
| Enrolled in school | 1282 | 727 | 56.7% | 1201 | 686 | 57.1% |
| In relationship | 1279 | 729 | 57.0% | 1201 | 685 | 57.0% |

**Predicting alcohol dependence symptoms with GPS above levels of consumption**

| Linear Mixed Models for AD Symptoms (N = 1,201) | | | | | | |
| --- | --- | --- | --- | --- | --- | --- |
|  | Model 1 | | | Model 2 | | |
|  | B | SE |  | B | SE |  |
| Drinking Frequency |  |  |  |  |  |  |
| Female | -0.213 | 0.059 | *** | -0.061 | 0.056 |  |
| Age | -0.001 | 0.040 |  | 0.005 | 0.038 |  |
| GPS | 0.123 | 0.030 | *** | 0.085 | 0.028 | ** |
| Drinking frequency | - | - | - | 0.505 | 0.040 | *** |
| Pseudo-R^2^ | 0.026 |  |  |  | 0.141 |  |
| Intoxication Frequency |  |  |  |  |  |  |
| Female | -0.213 | 0.059 | *** | 0.021 | 0.056 |  |
| Age | -0.001 | 0.040 |  | -0.009 | 0.036 |  |
| GPS | 0.123 | 0.030 | *** | 0.075 | 0.027 | ** |
| Intoxication frequency | - | - | - | 0.727 | 0.049 | *** |
| Pseudo-R^2^ | 0.026 |  |  |  | 0.182 |  |

Models include random intercepts for the family level. Pseudo-R2 calculated using the method described Nakagawa and Schielzeth (2013).

**Models with correction for confounding in GxE estimates (Keller, 2014)**

The following mixed models include interactions between both genome wide polygenic scores and relationship status, and each covariate (age, educational attainment, female, and currently enrolled in school).

| Linear Mixed Models for Drinking Frequency (N = 1,201) | | | | | | | | | |
| --- | --- | --- | --- | --- | --- | --- | --- | --- | --- |
|  | Model 1 | | | Model 2 | | | Model 3 | | |
|  | B | SE |  | B | SE |  | B | SE |  |
| Female | -0.456 | 0.059 | *** | -0.422 | 0.086 | *** | -1.344 | 1.290 | *** |
| Age | -0.003 | 0.041 |  | 0.070 | 0.058 |  | 0.070 | 0.058 |  |
| In relationship | -0.090 | 0.055 |  | 2.722 | 1.566 |  | 2.697 | 1.565 |  |
| GPS | 0.109 | 0.030 | *** | -2.408 | 0.837 | ** | -2.387 | 0.836 | ** |
| EDU: Basic | -0.248 | 0.123 | * | -0.341 | 0.192 |  | -0.340 | 0.192 |  |
| EDU: Secondary | 0.121 | 0.074 |  | 0.114 | 0.108 |  | 0.119 | 0.108 |  |
| EDU: Tertiary | -0.028 | 0.154 |  | -0.336 | 0.283 |  | -0.340 | 0.283 |  |
| In school | 0.055 | 0.066 |  | 0.069 | 0.096 |  | 0.064 | 0.096 |  |
| In relationship*GPS | - | - | - | -0.100 | 0.055 |  | -0.161 | 0.075 | ** |
| Female*In relationship | - | - | - | -0.040 | 0.112 |  | -0.042 | 0.112 |  |
| Age*In relationship | - | - | - | -0.128 | 0.071 |  | -0.127 | 0.071 |  |
| EDU: Basic*In relationship | - | - | - | 0.169 | 0.249 |  | 0.162 | 0.249 |  |
| EDU: Secondary *In relationship | - | - | - | 0.001 | 0.141 |  | -0.005 | 0.141 |  |
| EDU: Tertiary*In relationship | - | - | - | 0.308 | 0.332 |  | 0.311 | 0.332 |  |
| In school*In relationship | - | - | - | -0.017 | 0.127 |  | -0.009 | 0.127 |  |
| Female*GPS | - | - | - | 0.012 | 0.059 |  | -0.062 | 0.087 |  |
| Age*GPS | - | - | - | 0.115 | 0.038 | ** | 0.115 | 0.038 | ** |
| EDU: Basic *GPS | - | - | - | 0.139 | 0.120 |  | 0.150 | 0.120 |  |
| EDU: Secondary*GPS | - | - | - | 0.046 | 0.072 |  | 0.051 | 0.072 |  |
| EDU: Tertiary*GPS | - | - | - | -0.267 | 0.155 |  | -0.271 | 0.155 |  |
| In school*GPS | - | - | - | 0.040 | 0.066 |  | 0.035 | 0.066 |  |
| Female*In Relationship*GPS | - | - | - | - | - | - | 0.129 | 0.110 |  |
| Pseudo-R^2^ | 0.073 |  |  | 0.090 |  |  | 0.091 |  |  |

Models include random intercepts for the family level. Pseudo-R2 calculated using the method described Nakagawa and Schielzeth (2013).

| Linear Mixed Models for Intoxication Frequency (N = 1,201) | | | | | | | | | |
| --- | --- | --- | --- | --- | --- | --- | --- | --- | --- |
|  | Model 1 | | | Model 2 | | | Model 3 | | |
|  | B | SE |  | B | SE |  | B | SE |  |
| Female | -0.543 | 0.058 | *** | -0.534 | 0.084 | *** | -0.532 | 0.084 | *** |
| Age | 0.023 | 0.040 |  | 0.111 | 0.057 |  | 0.112 | 0.057 | * |
| In relationship | -0.178 | 0.054 | *** | 3.279 | 1.522 | * | 3.237 | 1.519 | * |
| GPS | 0.111 | 0.029 | *** | -1.764 | 0.818 | * | -1.727 | 0.817 | * |
| EDU: Basic | -0.289 | 0.120 | * | -0.406 | 0.187 | * | -0.405 | 0.186 | * |
| EDU: Secondary | 0.026 | 0.073 |  | -0.013 | 0.105 |  | 0.004 | 0.105 |  |
| EDU: Tertiary | -0.232 | 0.151 |  | -0.255 | 0.276 |  | -0.262 | 0.275 |  |
| In school | -0.094 | 0.065 |  | -0.067 | 0.093 |  | -0.076 | 0.093 |  |
| In relationship*GPS | - | - | - | -0.106 | 0.053 | * | -0.215 | 0.073 | ** |
| Female*In relationship | - | - | - | 0.002 | 0.109 |  | 0.002 | 0.109 |  |
| Age*In relationship | - | - | - | -0.158 | 0.069 | * | -0.157 | 0.069 | * |
| EDU: Basic*In relationship | - | - | - | 0.242 | 0.242 |  | 0.230 | 0.242 |  |
| EDU: Secondary *In relationship | - | - | - | 0.083 | 0.137 |  | 0.072 | 0.137 |  |
| EDU: Tertiary*In relationship | - | - | - | -0.038 | 0.323 |  | -0.030 | 0.322 |  |
| In school*In relationship | - | - | - | -0.078 | 0.123 |  | -0.063 | 0.123 |  |
| Female*GPS | - | - | - | -0.028 | 0.058 |  | -0.162 | 0.084 |  |
| Age*GPS | - | - | - | 0.084 | 0.037 | * | 0.084 | 0.037 | * |
| EDU: Basic *GPS | - | - | - | 0.029 | 0.117 |  | 0.049 | 0.117 |  |
| EDU: Secondary*GPS | - | - | - | 0.171 | 0.070 | * | 0.180 | 0.070 | * |
| EDU: Tertiary*GPS | - | - | - | -0.145 | 0.152 |  | -0.153 | 0.151 |  |
| In school*GPS | - | - | - | 0.032 | 0.064 |  | 0.024 | 0.064 |  |
| Female*In Relationship*GPS | - | - | - | - | - | - | 0.231 | 0.106 | * |
| Pseudo-R^2^ | 0.110 |  |  | 0.130 |  |  | 0.133 |  |  |

Models include random intercepts for the family level. Pseudo-R2 calculated using the method described Nakagawa and Schielzeth (2013).

| Linear Mixed Models for AD symptoms (N = 1,201) | | | | | | | | |
| --- | --- | --- | --- | --- | --- | --- | --- | --- |
|  | Model 1 | | | Model 2 | | | Model 3 | |
|  | B | SE |  | B | SE |  | B | SE |
| Female | -0.197 | 0.061 | *** | -0.125 | 0.089 |  | -0.124 | 0.089 |
| Age | -0.001 | 0.041 |  | 0.041 | 0.060 |  | 0.041 | 0.060 |
| In relationship | -0.097 | 0.057 |  | 1.370 | 1.642 |  | 1.355 | 1.642 |
| GPS | 0.123 | 0.030 | *** | 0.002 | 0.860 |  | 0.012 | 0.860 |
| EDU: Basic | 0.139 | 0.128 |  | 0.046 | 0.201 |  | 0.046 | 0.201 |
| EDU: Secondary | 0.033 | 0.076 |  | -0.052 | 0.112 |  | -0.050 | 0.112 |
| EDU: Tertiary | 0.079 | 0.160 |  | 0.037 | 0.296 |  | 0.035 | 0.295 |
| In school | -0.046 | 0.069 |  | 0.042 | 0.100 |  | 0.040 | 0.100 |
| In relationship*GPS | - | - | - | -0.112 | 0.057 |  | -0.141 | 0.079 |
| Female*In relationship | - | - | - | -0.126 | 0.117 |  | -0.127 | 0.117 |
| Age*In relationship | - | - | - | -0.065 | 0.074 |  | -0.064 | 0.074 |
| EDU: Basic*In relationship | - | - | - | 0.195 | 0.261 |  | 0.192 | 0.261 |
| EDU: Secondary *In relationship | - | - | - | 0.169 | 0.148 |  | 0.166 | 0.148 |
| EDU: Tertiary*In relationship | - | - | - | -0.054 | 0.348 |  | -0.053 | 0.348 |
| In school*In relationship | - | - | - | -0.169 | 0.133 |  | -0.165 | 0.133 |
| Female*GPS | - | - | - | 0.035 | 0.061 |  | 0.000 | 0.090 |
| Age*GPS | - | - | - | 0.007 | 0.039 |  | 0.008 | 0.039 |
| EDU: Basic *GPS | - | - | - | -0.129 | 0.125 |  | -0.123 | 0.125 |
| EDU: Secondary*GPS | - | - | - | 0.042 | 0.075 |  | 0.045 | 0.075 |
| EDU: Tertiary*GPS | - | - | - | -0.276 | 0.160 |  | -0.278 | 0.160 |
| In school*GPS | - | - | - | -0.003 | 0.068 |  | -0.005 | 0.068 |
| Female*In Relationship*GPS | - | - | - | - | - | - | 0.062 | 0.115 |
| Pseudo-R^2^ | 0.029 |  |  | 0.041 |  |  | 0.041 |  |

Models include random intercepts for the family level. Pseudo-R2 calculated using the method described Nakagawa and Schielzeth (2013).

**Levels of alcohol use/misuse across GPS Quintiles, Relationship Status, and Sex**

In order to assess whether outliers on the polygenic scores (near the high or low end of the distribution) were responsible for the GPS x relationship status interaction, we looked at observed values of each drinking phenotype across several scenarios. First, we collapsed individuals into quintiles of risk based on their GPS among those in a relationship and those not in a relationship. Next, we calculated the mean value of each alcohol phenotype (AD symptoms, drinking frequency, and intoxication frequency) and plotted them in the figure below. Finally, we repeated the process, amongst males and females to inspect for sex differences.

We see several patterns that help support the conclusions of our results. First, it is at the riskier end of the GPS distribution where differences in the alcohol use phenotypes between those in a relationship and those not in a relationship emerge. Second, it appears to be a monotonic increase across the risk continuum rather than extreme values among the riskiest category. Finally, we see that similar patterns across sex for AD symptoms, but different patterns across sex in terms of intoxication frequency (similar to results in the manuscript).

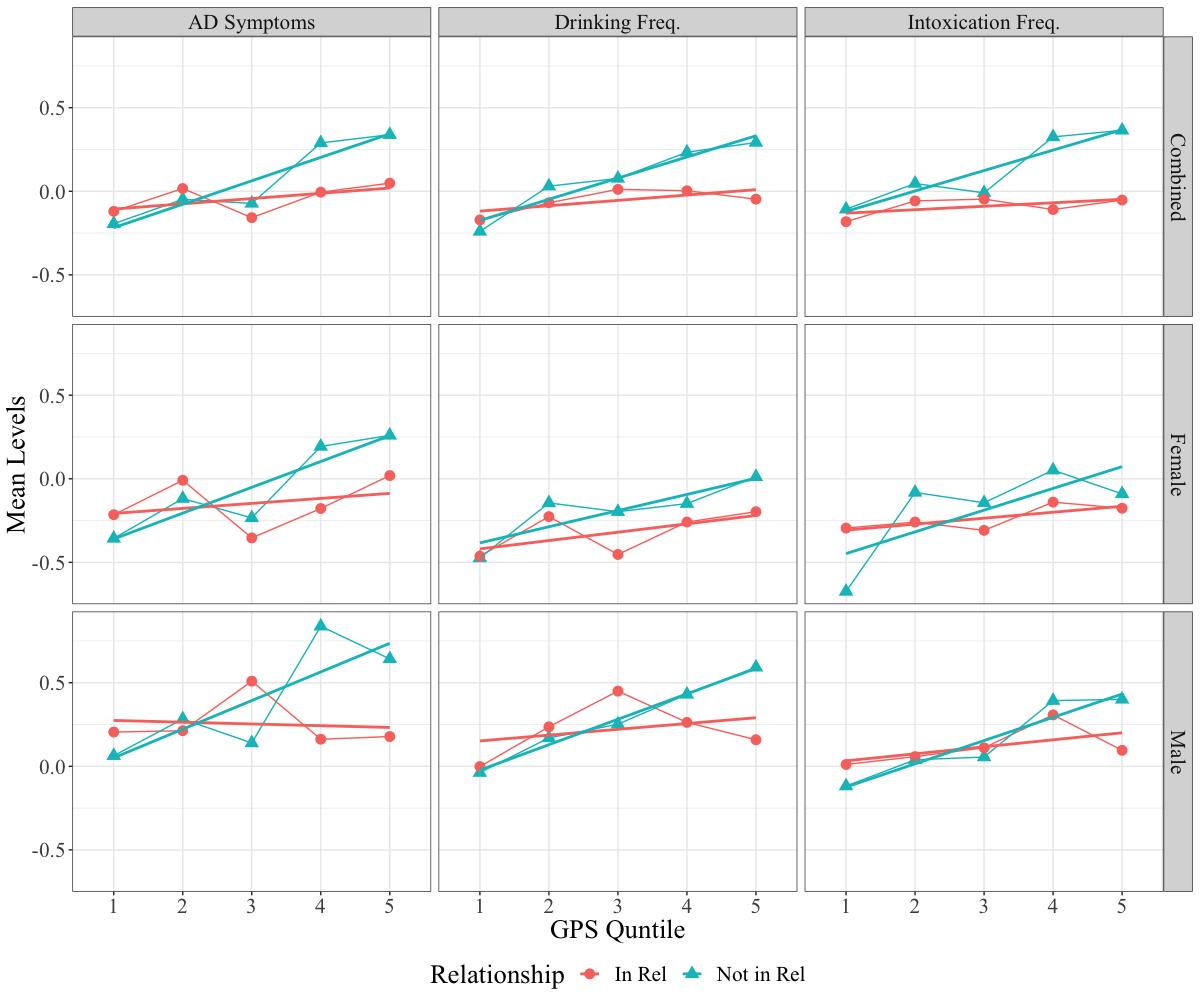

**Predictive Power of Alcohol-Related GPS**

We tested genome-wide polygenic scores derived from other alcohol related GWAS to determine whether other GPS would perform better for assessing genetic liability towards alcohol misuse. Because the discovery phenotype for the GSCAN^1^ GWAS was drinks per week (GSCAN DPW) measuring alcohol consumption) and measures of consumption and problems have related but distinct genetic influences^2,3^, we also tested polygenic scores derived from recent GWAS on the Alcohol Use Disorders Identification Test (AUDIT)^4^ and from the Psychiatric Genomics Consortium’s most recent GWAS of alcohol dependence (PGC AD)^5^. In using the AUDIT summary statistics, we further derived GPS specific to measures of consumption (AUDIT-C) and problems (AUDIT-P).

The figure below shows the incremental R^2^ (GPS above model with age and sex as covariates) for each phenotype with each GPS across a range of p-value thresholds. Regardless of the phenotype or threshold, the GPS derived from GSCAN were more predictive. This is likely the result of the significantly larger sample size in GSCAN compared to the other discovery GWAS (~1 million, compared to 121,604 in the AUDIT and 28,757 in the within-ancestry, unrelated sample for the PGC AD). Predictive power of GPS is driven primarily by sample size in the discovery GWAS^6^.

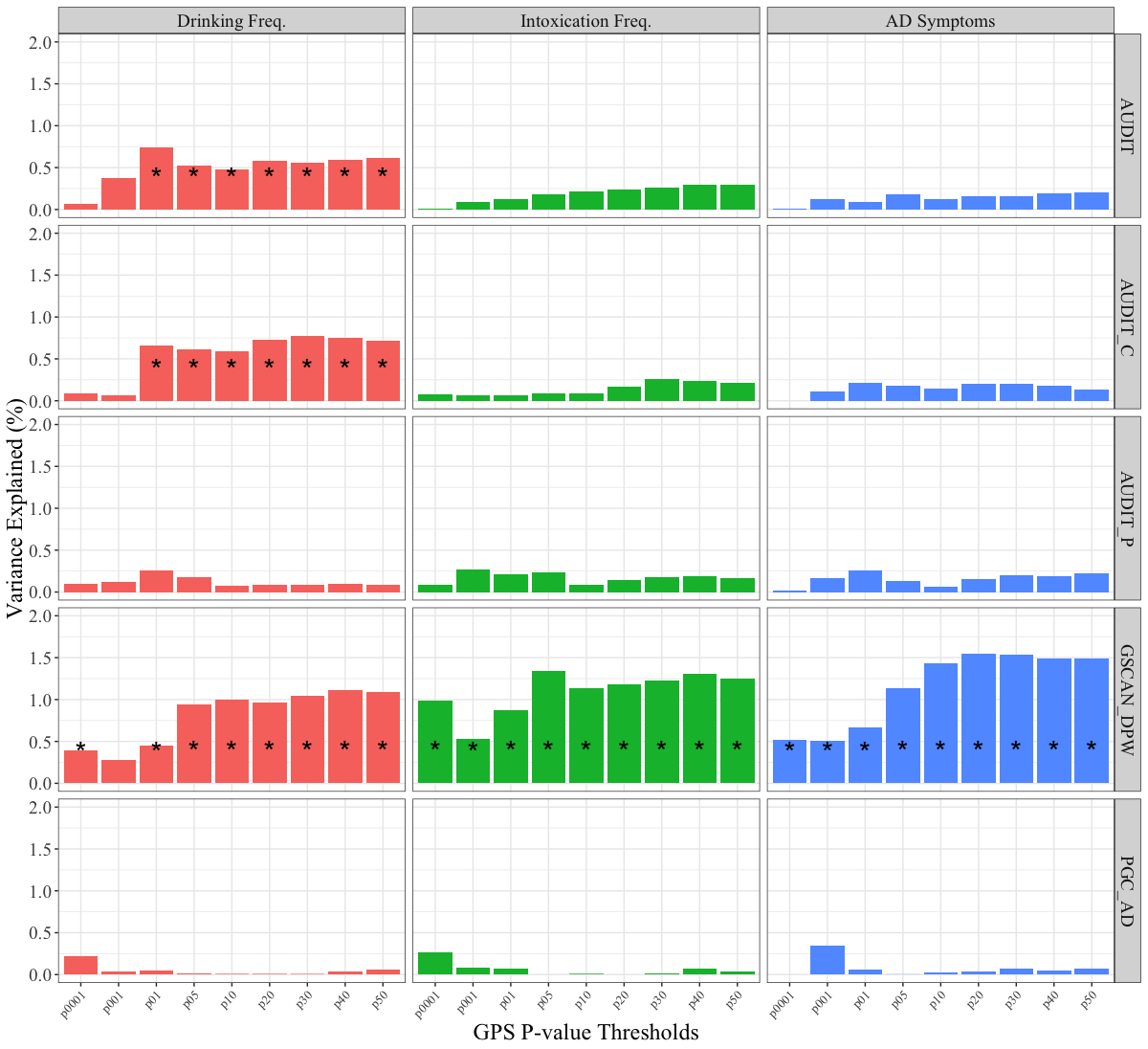

**Comparing the GxE effect between 12-month and Lifetime AD Symptom counts**

Our measure of DSM-IV symptom counts in the primary analyses included lifetime prevalence of symptoms. In order to determine whether the onset of symptoms predated relationship status, we ran a separate series of analyses with a dummy code for experiencing symptoms within the past year compared to those who have experienced symptoms more than 12 months prior (N= 165 in past year)

The table below shows the correlations among alcohol phenotypes. Regardless of whether AD symptoms were experienced in the previous 12 months or before that, they showed the same correlation with current drinking and intoxication frequency

Correlations Among Alcohol Phenotypes in FinTwin12

|  |  | 1 | 2 | 3 | 4 | 5 |
| --- | --- | --- | --- | --- | --- | --- |
| 1 | Drinking Frequency | 1.00 | - | - | - | - |
| 2 | Intoxication Frequency | 0.64 | 1.00 | - | - | - |
| 3 | AD symptoms (lifetime) | 0.33 | 0.41 | 1.00 | - | - |
| 4 | AD symptoms (≤ 12 months prior) | 0.20 | 0.24 | 0.43 | 1.00 | - |
| 5 | AD symptoms (> 12 months prior) | 0.20 | 0.26 | 0.74 | -0.28 | 1.00 |

The figure below plots the results from linear mixed models containing an additional indicator for whether or not symptoms were experienced in the past 12 months compared to those experienced earlier. Model 1 contains only the main effect of this dummy, while Model 2 includes the interaction between this dummy and relationship status, GPS, and the relationship*GPS interaction term.

Including the interactions (Model 2) did not result in a better fitting model (Δ-2LL = 0.452, df = 3, p = 0.929), suggesting that early and later onset of AD symptoms only differed in their intercepts rather than the slopes for these variables (e.g. Model 1 is the better fitting model). Though the confidence intervals overlap in the 12-month category, the slopes are in the same direction as the previous GxE results with lifetime AD symptoms. Additionally, there are a smaller number of people reporting past year symptoms relative to those reporting earlier (N= 165 report any symptoms in the past year, N= 666 report any symptoms prior to the past year).

As both earlier and current symptoms are equally correlated with the other alcohol phenotypes, and the pattern of the GxE interaction is the same across these groups, the interaction for AD symptoms in the manuscript most likely reflects moderation of the shared variance between AD symptoms and current intoxication frequency.

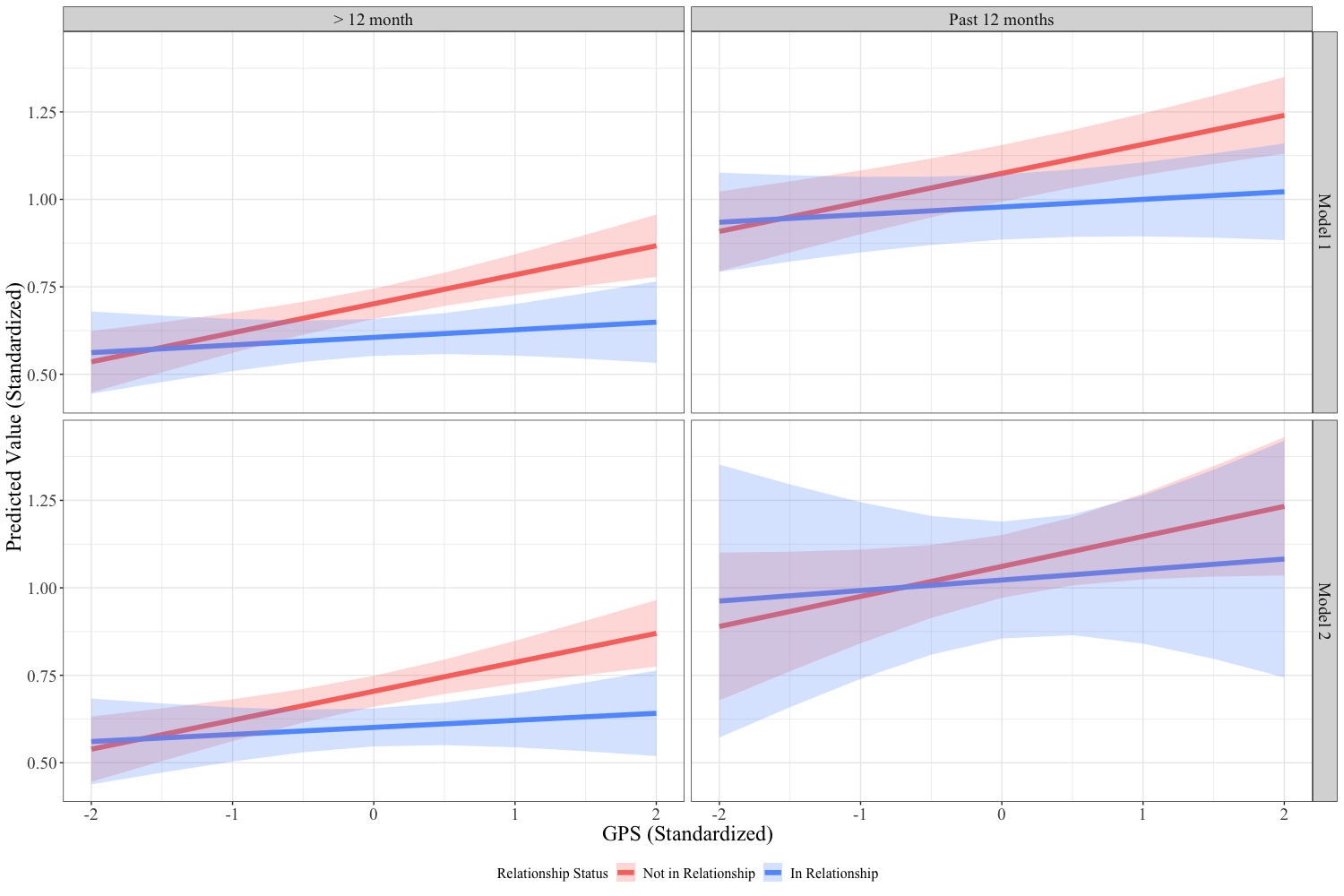

**Comparison of Between and Within Family Effects**

Because difference between families (e.g. neighborhood factors, religiosity, socioeconomic status, population stratification) can confound any of the observed relationships between 1) relationship status, 2) genome-wide polygenic scores, or 3) their interaction and each of the alcohol phenotypes, it is important to utilize methods that can adjust for this confounding. One common approach is to utilize family fixed-effects (within-family analyses) to adjust for any between family confounders. The logic of this type of model is to examine whether differences between siblings in an exposure/risk factor relate to differences in alcohol misuse. Thus, in the equation^7^:

*Y_ij_ = βX_ij_ + γW_i_ + α_i_ + ε_ij_*

the effect of the vector of within-family risk factors *X* on *Y* for twin *i* in family *j* is conditional upon a vector of covariates that vary between family (e.g., socioeconomic status, parental alcohol abuse, religiosity), *W*, and another vector of unmeasured fixed effects that vary between family, *α*, plus the family specific random error term, *ε_ij_*. If we were to imagine this as a comparison of two siblings, the equation could be expressed as:

*Y_i2_ - Y_i1_ = βX_i2_ + γW_i_ + α_i_ + ε_i2_* – (*βX_i1_ + γW_i_ + α_i_ + ε_i1_*)*= β*(*X_i2_ - X_i1_*) *+* (*ε_i2_* –*ε_i1_*).

The effects of all covariates that do not vary within families are therefore cancelled out of the model. The distinct inferential advantage of a twin fixed effects design is that it allows us to examine whether predictors are robust after controlling for familial confounding (i.e., all of the genetic and shared environmental factors that twin siblings share), enabling stronger inferences not possible in samples of singletons. We represent the result from the within-family (fixed effects) compared to between family (random effects) estimates in Relationship Status, GSCAN GPS, and their interaction graphically in the figure below. We limited our sample to DZ twins (N=699 twins from 405 families), as we are also interested in the difference in GPS (which does not vary between MZ twins). We see similar, though attenuated effects in the within family models. This could be the result of confounding in the between family estimates as well as reduced power in the within family models.

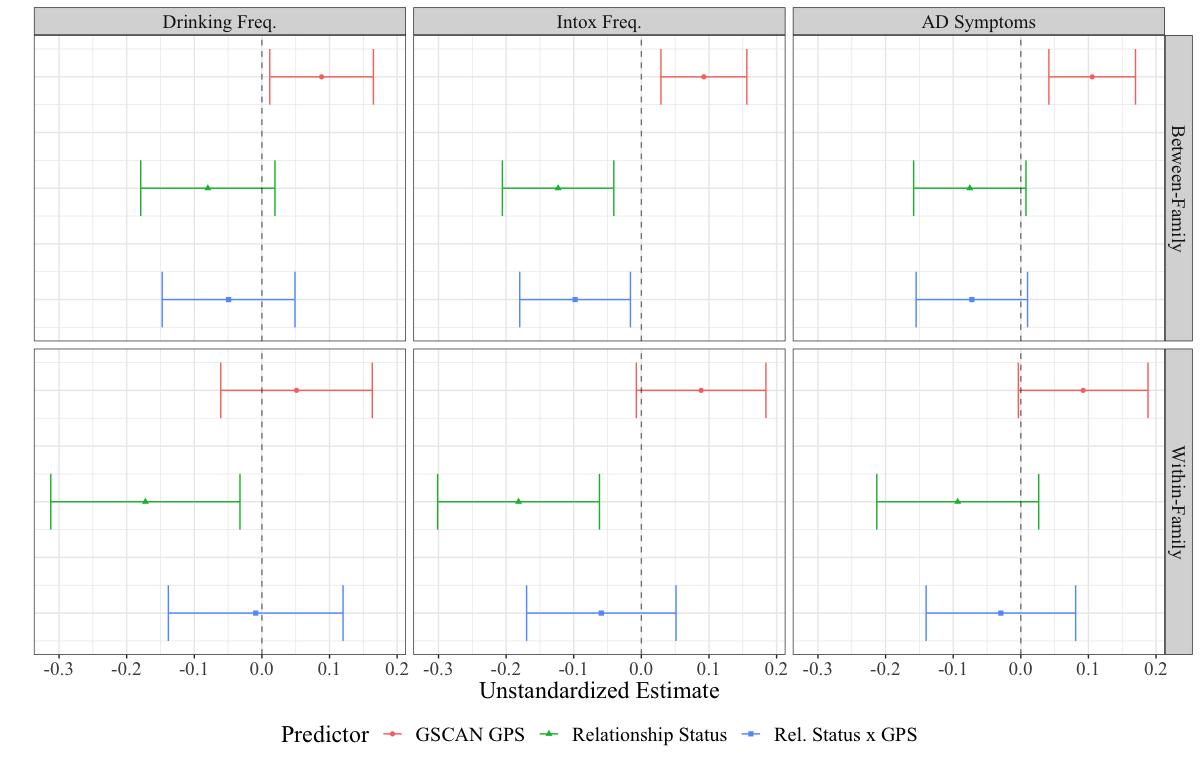
